# Evaluating drone-mounted thermal infrared sensors for macropod monitoring in Tasmania

**DOI:** 10.1101/2025.11.27.691070

**Authors:** Yee Von Teo, Darren Turner, Jessie C. Buettel, Barry W. Brook

## Abstract

Thermal drones offer significant advantages for monitoring wildlife in low-light conditions; however, detection performance is influenced by technical settings and environmental factors. This study evaluates the use of nocturnal drone surveys to detect macropods in Tasmania, with the aim of determining optimal flight parameters for balancing detection accuracy and survey efficiency. Field surveys were conducted in Narawntapu National Park using varying flight altitudes and image overlap settings, and detection rates were compared across thermal datasets. Manual annotations and basic thresholding method were used to quantify detection success. Results showed that detection rates were highest when surveys were conducted under cooler ambient temperatures and at moderate altitudes (e.g. 60 metres AGL) with 50% image overlap. These findings provide practical guidance for designing nocturnal drone surveys and offers baseline recommendations for using drone-mounted thermal sensors to monitor large-bodied, crepuscular mammals, with broader implications for scalable wildlife monitoring programs.

## 1 Introduction

Wildlife monitoring is crucial for effective conservation and management. Yet nocturnal and crepuscular species can be especially challenging to observe, owing to their peak activity in low-light conditions and their elusive behaviours (Hart *et al.* 2022). Traditional ground-based techniques, such as camera trapping (Tan, Yang & Niu 2013; Caravaggi *et al.* 2018) and spotlighting (Ruette, Stahl & Albaret 2003; Wayne *et al.* 2005; Sanders *et al.* 2024), are commonly employed to estimate population size, assess relative abundance, or determine occupancy of these species. While these methods are widely used and can be highly effective in certain contexts, they present notable limitations, and certain objectives, such as detecting rare or highly mobile animals, can be difficult to achieve. Camera traps have a limited detection zone, capturing only animals within a narrow field of view (Zwerts *et al.* 2021; Palencia *et al.* 2022). Spotlight surveys, although more mobile, are time-intensive and susceptible to observer bias, differences in operator skill, and unpredictable animal behaviour (Ruette, Stahl & Albaret 2003; Sunde & Jessen 2013). A particularly challenging aspect of both methods is the reduced detectability of target species, particularly in areas with dense vegetation or obstructive environmental features.

In response, some studies have explored handheld or vehicle-mounted thermal imagers to detect nocturnal wildlife along transects (Focardi 2001; Augusteyn, Pople & Rich 2020; McGregor *et al.* 2021; Pocknee *et al.* 2021; Underwood, Derhè & Jacups 2022). For instance, Underwood, Derhè and Jacups (2022) found that thermal imaging surveys using a handheld imager increased detection rates and successfully located species often missed by spotlighting in rainforest habitats. Similarly, McGregor *et al.* (2021) reported improved detection of small-and medium-sized mammals using vehicle-based thermal imaging in open, arid environments. However, like spotlight surveys, ground-based thermal detection is limited by a narrow field of view and often requires slow or repeated passes to ensure comprehensive coverage, and are typically constrained to accessible roads or tracks. This reduces their applicability in remote or rugged terrain.

Researchers have also explored other efficient and objective alternatives. One early example of remote sensing for nocturnal wildlife detection was proposed in the late 1960s, when researchers mounted an infrared line scanner on a crewed aircraft (Croon *et al.* 1968) to detect white-tailed deer (*Odocoileus virginianus*). Despite the innovation of this approach, this early work faced several technological constraints: thermal systems were prohibitively expensive at the time, and detection success was limited in areas with dense forest canopy that obstructed the line of sight to animals on the ground.

With advances in drone platforms and miniaturised thermal sensors, nocturnal drone surveys are now a viable and increasingly accessible option for wildlife monitoring (Mirka *et al.* 2022; Whitworth *et al.* 2022; Santos *et al.* 2023; Wijayanto, Condro & Rahman 2023). Over the past decade, these surveys have shown considerable promise in detecting endothermic species across a range of habitats and environmental conditions (Rahman *et al.* 2020; Tovar-Sánchez *et al.* 2021; Sellés-Ríos *et al.* 2022; Larsen *et al.* 2023; Olsen, Barron & Butler 2023; Trinh-Dinh *et al.* 2024). Drone-based thermal surveys offer a unique aerial perspective and can operate in low-light conditions, providing a powerful alternative to traditional ground-based methods.

Macropods, including kangaroos and wallabies, are relatively large marsupials that often inhabit open or semi-open habitats and play important roles in ecosystem functioning as grazers and seed dispersers (Ramsey & Wilson 1997; Southwell *et al.* 1999; Chapman 2003; Wiggins *et al.* 2010). Their size and habitat preferences make them especially amenable to detection with aerial thermal imagery. Yet despite the growing interest in thermal drone applications, few studies have focused specifically on their application to macropod detection (Gentle *et al.* 2018; Brunton, Leon & Burnett 2020), and many questions remain regarding how to optimise flight operations for effective, low-disturbance detection. For example, Brunton, Leon and Burnett (2020) examined different flight altitudes for monitoring Eastern grey kangaroos (*Macropus giganteus*) across various habitats in Queensland, but broader investigations into how flight parameters influence survey outcomes remain limited. Comparable studies on other large-bodied mammals have begun to explore similar trade-offs (Rahman *et al.* 2020; Kim, Chung & Lee 2021; Larsen *et al.* 2023), but few studies have systematically evaluate flight parameters—including flight altitude and image overlap percentage—specifically for macropods in Tasmania, representing a clear gap in the literature.

This study addresses that gap by evaluating the feasibility of drone-based thermal imaging for detecting macropods under low-light conditions. The specific objectives of this study were to: (1) identify the optimal flight parameters—particularly image overlap percentage and flight altitude—that maximise detection probability while ensuring comprehensive coverage of the survey area, since greater overlap reduces the chance of missed detections and supports post-processing; and (2) examine the suitability of manual annotation, as compared with a simple image thresholding approach, for detecting macropods in thermal imagery. Additionally, a preliminary application of image thresholding is presented to compare the time efficiency of manual and automated detection workflows.

## 2 Materials and Methods

### 2.1 Study sites

This study was conducted at Narawntapu National Park (Latitude: –41.149141, Longitude: 146.603217), located on the central north coast of Tasmania, Australia (Figure 1). This park features a mosaic of coastal heathlands, wetlands, dry sclerophyll woodlands, and open grasslands, providing key habitat for native marsupials (Parks and Wildlife Service 2016; Martin *et al.* 2018; Teo *et al.* 2024). Historically, parts of the park were grazed, leaving open areas interspersed with regenerating vegetation. These modified yet structurally open habitats continue to support a high density of herbivorous marsupials (Parks and Wildlife Service 2016), particularly Bennett’s wallabies (*Notamacropus rufogriseus*) and Tasmanian pademelons (*Thylogale billardierii*). The number of the Forester kangaroo (*Macropus giganteus tasmaniensis*) declined in 2019 but reportedly reached approximately 60 individuals per kilometre in 2020 (Tamura *et al.* 2021).

**Figure 1.**
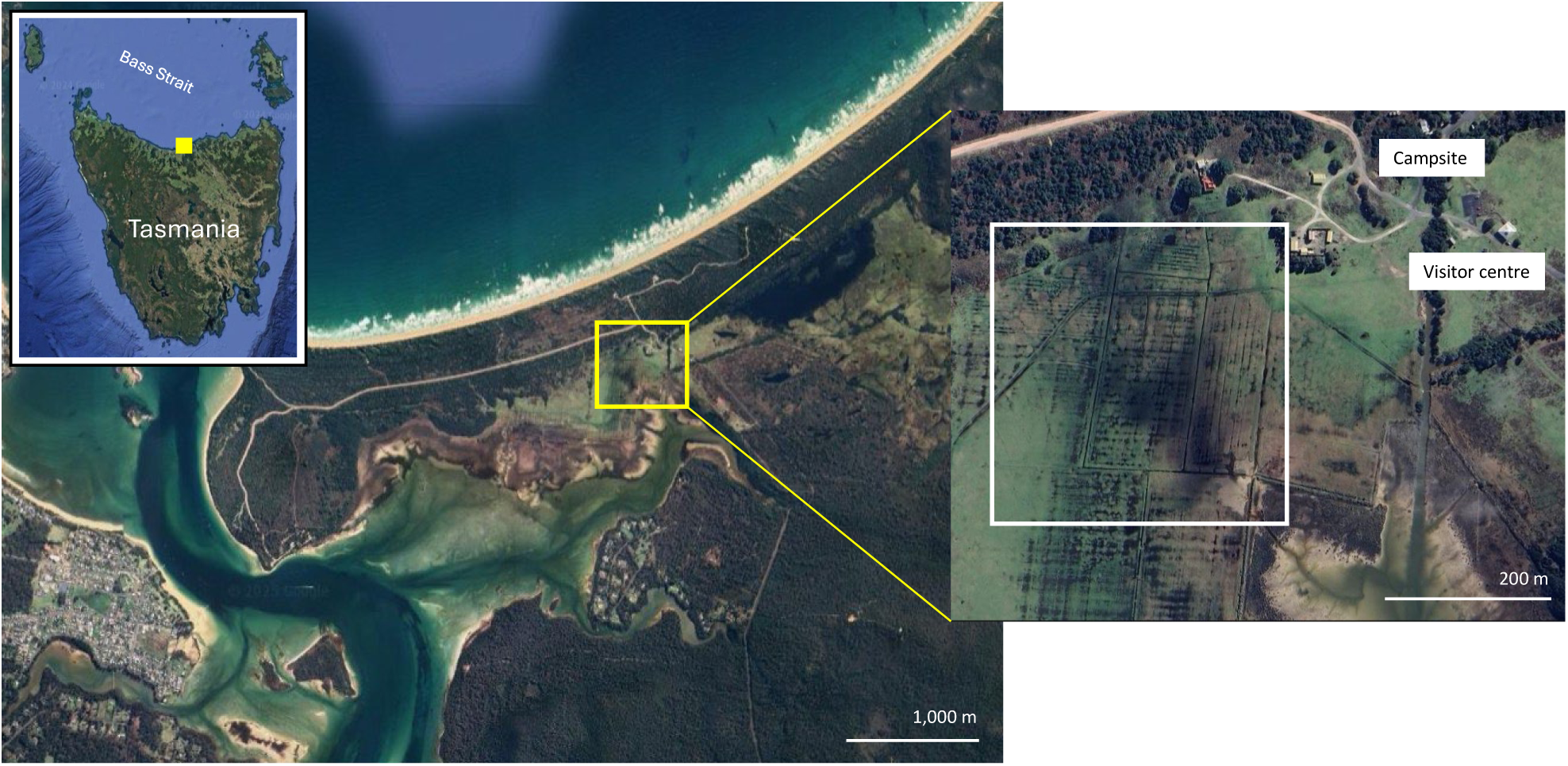
Map of the study area within Narawntapu National Park, Tasmania. The inset map highlights the specific survey area (white box) where field data collection was conducted.

Although all three species co-occurred across the survey area, Bennett’s wallabies and Forester kangaroos were commonly observed in open, grassy areas, while pademelons were favoured woodland edges or denser vegetation (Le Mar & McArthur 2005). Forester kangaroos are relatively large macropods (Tanner & Hocking 2001), whereas Bennett’s wallabies and pademelons are smaller-bodied (Le Mar, Southwell & McArthur 2001). These differences in body size and spatial use may influence thermal detectability, especially in areas with variable vegetation density or partial occlusion, as smaller species like pademelons may be more easily obscured in dense habitats compared to larger kangaroos in open grasslands.

The selected survey area is an open grassland adjacent to low bushland, chosen for its frequent macropod activity and minimal human disturbance. Located away from visitor facilities, it minimises anthropogenic interference during surveys. The target species present at the site included the Forester kangaroo, Bennett’s wallaby, and the Tasmanian pademelon. These species are commonly observed grazing in the open grasslands or near the edges of the low bushland, particularly during crepuscular periods, and their frequent presence made the site suitable for evaluating the effectiveness of drone-based thermal detection methods.

### 2.2 Drone platform and thermal sensor

We deployed a DJI Mavic 3 Thermal quadcopter, outfitted with a visual (RGB) camera and a thermal sensor. The RGB camera offers a maximum image resolution of 8000 × 6000 pixels, facilitated by a 1/2’’ CMOS sensor with 48 effective megapixels. The thermal imaging component was an uncooled VOx (vanadium oxide) microbolometer with a resolution of 640 × 512 pixels and a frame rate of 30 Hz. The Mavic 3T has a maximum flight time of approximately 45 minutes under ideal conditions (no wind or very low wind speed) without additional payload.

### 2.3 Flight operations

We carried out nocturnal drone surveys under stable weather conditions with wind speeds below 20 km/h to optimise flight stability and minimise battery consumption. Flight missions were carried out during the early hours of the day between 0000 and 0700 local time. To minimise disturbance to the target species, the drone was launched at a minimum distance of 150 metres from the nearest visible animal. All flights were pre-programmed in the DJI Pilot 2 application using a standard lawnmower (grid) pattern for systematic coverage. The gimbal was set to a nadir orientation, pitched at –90°, to ensure the thermal sensor remained perpendicular to the ground during data collection.

Surveys were conducted at three flight altitudes: 40 m, 60 m, and 80 m above ground level (AGL). The minimum altitude of 40 metres AGL was selected based on species-specific protocols developed in a previous study (Teo *et al.* 2024), which aimed to minimise disturbance to the target macropod species. Detection success was found to decline significantly above 80 metres AGL, with thermal signatures becoming indistinct at higher elevations. Therefore, 80 metres AGL was used as the maximum operational ceiling. To reduce image blur and maintain adequate thermal resolution, flight speed was adjusted according to altitude: the drone was flown at 2.5 ms⁻¹ at 40 metres AGL, 3.75 ms⁻¹ at 60 metres AGL, and 5.0 ms⁻¹ at 80 metres AGL. These speeds were selected empirically during preliminary flights to balance image sharpness and coverage efficiency, with slower speeds at lower altitudes preserving detail for orthomosaic reconstruction, and faster speeds acceptable at higher altitudes due to the wider field of view.

At each altitude, front and side image overlaps of 70/80%, 50/50%, and 20/20% were tested to explore how overlap affects detection and orthomosaic accuracy. Higher % overlap provide more comprehensive coverage and is essential for generating high-quality orthomosaics; lower overlaps reduce the chance of double counting moving animals but potentially miss individuals at the edge of frames. This variation allowed for evaluating the trade-offs between coverage quality and detection reliability.

Furthermore, two different area extents: a smaller focal area (A1) and a larger area coverage (A2), were surveyed at each combination of altitude and overlap setting. Both coverage sites were located within the same broader study area to ensure environmental consistency. The comparison of small versus large areas allows assessment of scale-dependent detection patterns, i.e., whether detection probability or efficiency differs when surveying a concentrated, focal area versus a more extensive area with the same flight parameters. A third area (A3) close to Site A1 and A2 was also surveyed to provide supplementary data.

#### 2.3.1 Environmental conditions

All flights took place between 0000 and 0700 h, so the daily minimum temperatures likely reflect conditions during these surveys. To account for the residual ground heat influencing thermal contrast, maximum temperatures from the previous day of the flight operations were recorded. The daily global solar exposure, which is the total amount of solar energy that reaches a horizontal surface in a day, was also shown in Table 1. All temperature and solar exposure data were obtained from the Bureau of Meteorology (BoM) website (Bureau of Meteorology 2025).

**Table 1.** Weather conditions for each survey night based on BoM records (Bureau of Meteorology 2025). Maximum temperature and solar exposure values are taken from the preceding day, as these influence residual ground heat during nocturnal thermal surveys.

| Survey date | Max temp from previous day (°C) | Min temp (°C) | Solar exposure (MJ/m<sup>2</sup>) |
| --- | --- | --- | --- |
| 30<sup>th</sup> Sep | 19.7 (29<sup>th</sup> Sep) | 11.8 | 19.5 (29<sup>th</sup> Sep) |
| 1<sup>st</sup> Oct | 16.7 (30<sup>th</sup> Sep) | 9.6 | 13.2 (30<sup>th</sup> Sep) |
| 2<sup>nd</sup> Oct | 19.7 (1<sup>st</sup> Oct) | 3.9 | 18.6 (1<sup>st</sup> Oct) |

### 2.4 Image processing

Agisoft Metashape Professional was used to generate orthomosaics from each flight. However, 20/20% overlap was insufficient for complete alignment, likely due to the reduced contrast and increased noise typical of thermal imagery captured at night. Unlike RGB data, thermal imagery requires more overlap and distinct features to generate enough tie points for successful stitching (Cui *et al.* 2021). Additionally, one image set captured at 80 metres AGL with 50% front and side overlap also failed to align and was excluded from further analysis. Even when successfully aligned, those with lower overlap settings (e.g. 20/20%) appeared visually patchy, with noticeable gaps between stitched images, making them less suitable for accurate annotation and analysis. The size of each orthomosaic was calculated in hectares (ha).

#### 2.4.1 Manual annotation

All successfully generated orthomosaics were manually annotated to locate individual macropods, and annotation time was recorded. All three species present in the study area were labelled collectively as “macropods” rather than by species. Orthomosaics from flights with insufficient overlap were excluded from manual annotation due to incomplete image coverage.

#### 2.4.2 Thresholding

In parallel, a pixel-based thresholding approach was tested on each orthomosaic to automate macropod detection. Thresholding on thermal imagery provided a simple, rapid, and effective proof-of-concept workflow, demonstrating that the detection methodology is robust even with a straightforward approach and providing a solid foundation for future, more advanced computer vision applications, such as Convolutional Neural Networks (CNNs), to automate and extend detection capabilities. Image processing using thresholding was done in Python

3.7.1 with an 11th Gen Intel® Core™ i9-11900 CPU @ 2.50GHz (8 cores, 16 threads) and 128 GB of RAM. No dedicated GPU was used. A custom Python function was developed using

OpenCV to detect localised heat signatures (‘hotspots’). Intensity-based thresholding, morphological filtering, and local contrast evaluation were combined to isolate regions resembling animal targets.

The detection algorithm involved the following key steps:

1. Greyscale conversion: The thermal images, stored as JPEGs, were converted to 8-bit greyscale intensity, 𝐼𝐼_𝑔𝑔𝑔𝑔𝑔𝑔𝑔𝑔_, to simplify intensity analysis based on the following equation:

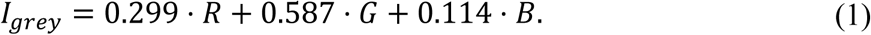

The coefficients in Equation (1) are defined by the ITU-R BT.601 standard (OpenCV 2024). This step was used to reduce image dimensionality and enhance intensity-based contrast.

1. Thermal thresholding: A relative threshold, 𝑇𝑇, was applied to isolate pixels with high thermal intensity according to Equation (2):

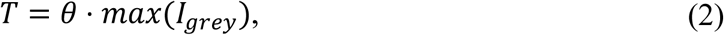

where 𝜃 is the heat threshold. Pixels with 𝐼𝐼_𝑔𝑔𝑔𝑔𝑔𝑔𝑔𝑔_ ≥ 𝑇𝑇 were retained in a binary mask, and contiguous bright regions were extracted as contours.

1. Geometric filtering: Each detected contour was evaluated based on its area 𝐴𝐴, which was given by the number of pixels, and aspect ratio 𝐴𝐴𝑅𝑅 = <u>^𝑤𝑤^</u>, where 𝑤𝑤 and ℎ are the width and height of the bounding box, respectively. Only contours satisfying:

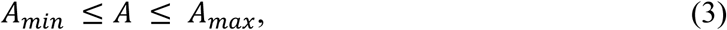

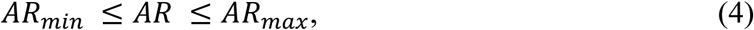

were retained. The allowable aspect ratio range was bounded by the reciprocal and multiple of a maximum ratio (e.g., 1/5 to 5). These thresholds were informed by visual inspection of annotated images. Values outside these bounds typically reflected background artefacts or thermal noise. The area range captured the approximate pixel dimensions of macropods across flight altitudes, while the aspect ratio range accommodated typical body elongation in various poses (e.g., standing, lying, or partially obscured).

1. Local Contrast Evaluation: To minimise false positives from uniformly bright areas (e.g., rocks or sunlit vegetation), a local contrast check was performed. For each candidate region, contrast was calculated using Equation (5).

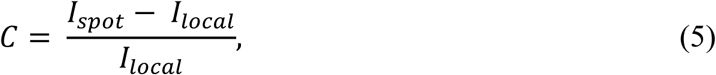

where 𝐼𝐼_𝑠𝑠𝑠𝑠𝑠𝑠𝑠𝑠_is the mean greyscale intensity of the hotspot, 𝐼𝐼_𝑙𝑙𝑠𝑠𝑙𝑙𝑚𝑚𝑙𝑙_is the mean intensity of surrounding pixels within a defined radius (excluding the hotspot itself). Only regions in which 𝐶𝐶 ≥ 𝛿𝛿, where 𝛿𝛿 is the local contrast threshold, were considered valid hotspots.

Because flights were flown across multiple nights and under varying ambient temperatures, the thermal contrast between macropods and the background varied between orthomosaics, requiring adjustment to the thresholding parameters (Table 2). For image sets classified as ‘warm’ (with higher background heat), a heat threshold of 0.7 and local contrast of 0.4 were applied. For datasets with colder surroundings, these values were reduced to 0.5 and 0.3 respectively. These parameter values were selected through preliminary trials on image subsets from both warm and cold conditions to maximise contrast between animal signatures and the background, while minimising false positives from residual ground heat or noise.

**Table 2.**
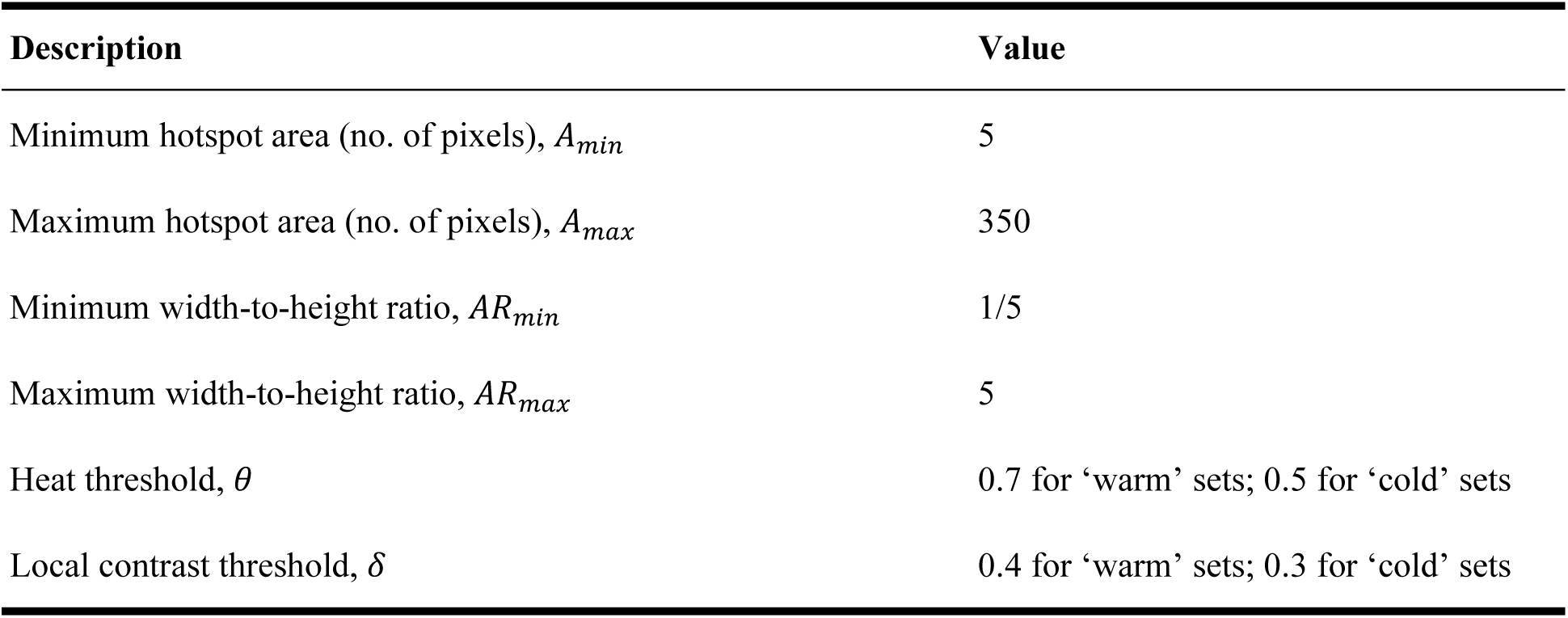
Parameter values and descriptions for the automated thresholding method implemented for macropod detection.

#### 2.4.3 Performance metrics

The performance of the thresholding method was evaluated using the commonly applied detection metrics, which include precision, recall, and F₁ score. A detected hotspot was treated as a true positive if it corresponded spatially to an annotated macropod in the orthomosaic; otherwise, it was classified as a false positive. Annotated macropods with no corresponding bounding box were considered false negatives. 𝑃 quantifies the proportion of detected bounding boxes that corresponded to true macropod detections and is defined as Equation (6). High precision indicates that macropod locations were accurately identified in the orthomosaic, with few false positives.

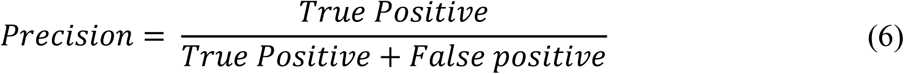

measures the proportion of actual objects (i.e., macropods) in the orthomosaic that have been correctly detected, as shown in Equation (7). High recall indicates that the performance of the thresholding method is effective at finding and localising the objects, with very few being missed (i.e., minimal false negatives).

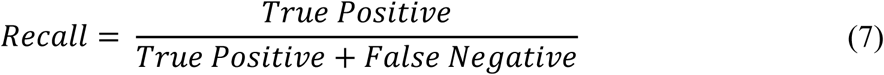

Overall performance was summarised using the 𝐹𝐹_1_score, calculated for each orthomosaic as the harmonic mean of precision and recall. The F₁ score ranges from 0 to 1, where 1 indicates perfect precision and recall.

The F₁ score is defined by Equation (8):

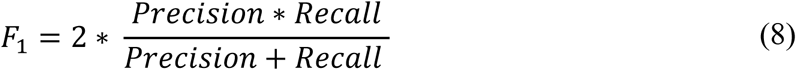

## 3 Results

### 3.1 Flight parameters

This section presents results relating to the drone flight parameters tested during nocturnal drone surveys, including flight altitude and image overlap. These parameters influenced key survey outcomes such as spatial coverage, flight duration, and overall operational efficiency.

The following subsections detail these outcomes and highlight the trade-offs between resolution, coverage, and flight duration.

#### 3.1.1 Spatial Coverage and Flight Duration

Flight altitude and image overlap percentage had significant impacts on both spatial coverage and flight duration. As shown in Table 3, increasing flight altitude from 40 metres to 80 metres AGL at 70/80% overlap reduced flight duration from 122 minutes to 27 minutes at Site A2, while increasing coverage from 19.71 ha to 21.59 ha (Table 4). Decreasing overlap percentage from 70/80% to 50/50% substantially reduced flight time while maintaining similar coverage areas (Figure 2).

**Figure 2.**
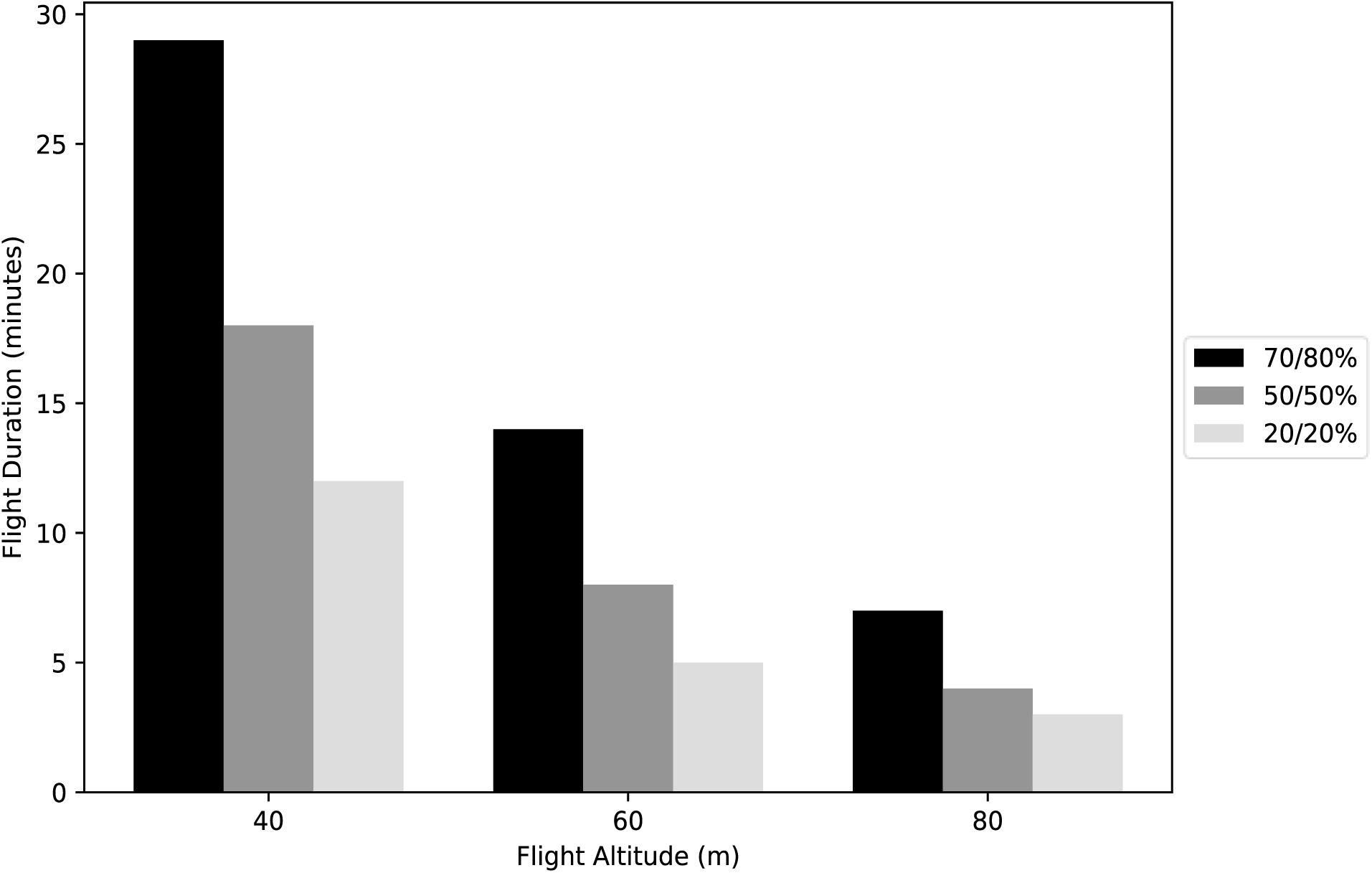
Flight duration at varying altitudes, with overlap percentage indicated in the legend. The bar chart demonstrates a trend of decreasing flight duration with increasing altitude. Data presented are limited to Site A1.

**Table 3.**
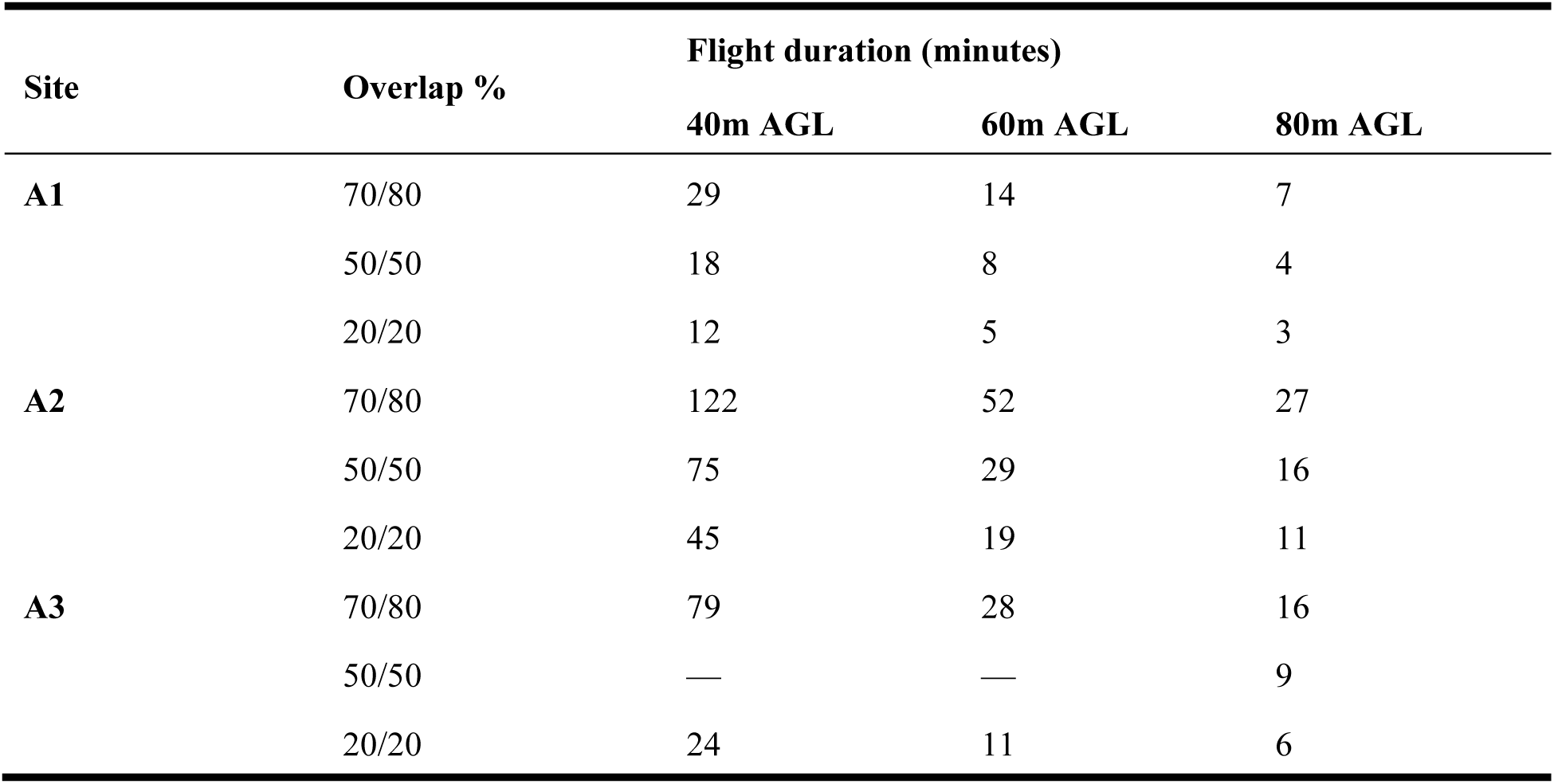
Flight duration (minutes) for three study sites (A1, A2, A3) across varying flight altitudes (40 m, 60 m, and 80 m above ground level: AGL) and image overlap percentages (70/80%, 50/50%, and 20/20%). Higher flight altitudes and lower overlap percentages consistently reduced survey time across all sites. Missing data (denoted by ’—’) reflects incomplete data collection at Site A3, which was designated as a secondary study location.

**Table 4.** Spatial coverage (hectares) for three study sites (A1, A2, A3) across varying flight altitudes (40 m, 60 m, and 80 m above ground level: AGL) and image overlap percentages (70/80%, 50/50%, and 20/20%). Asterisks (*) indicate parameter combinations where orthomosaic generation failed due to insufficient image alignment. Missing data (denoted by ’—’) reflects incomplete data collection at Site A3, which was designated as a secondary study location.

| Site | Overlap % | Spatial coverage (ha) |  |  |
| --- | --- | --- | --- | --- |
|  |  | 40m AGL | 60m AGL | 80m AGL |
| A1 | 70/80 | 6.2 | 6.71 | 6.65 |
|  | 50/50 | 5.24 | 5.44 | 5.67 |
|  | 20/20 | * | * | * |
| A2 | 70/80 | 19.71 | 21.51 | 21.59 |
|  | 50/50 | 18.67 | 20 | * |
|  | 20/20 | * | * | * |
| A3 | 70/80 | 12.07 | 13.41 | 15.78 |
|  | 50/50 | — | — | 11.83 |
|  | 20/20 | * | * | * |

However, overlap combinations of 20/20% frequently failed to produce usable orthomosaics, resulting in major gaps in spatial coverage. This trend was consistent across all sites (Table 4), although the figure presentations in this section focus on Site A1 due to its complete data coverage (Figure 3). Similar coverage and duration patterns were observed at Sites A2 and A3, where low-overlap combinations again led to alignment failures.

**Figure 3.**
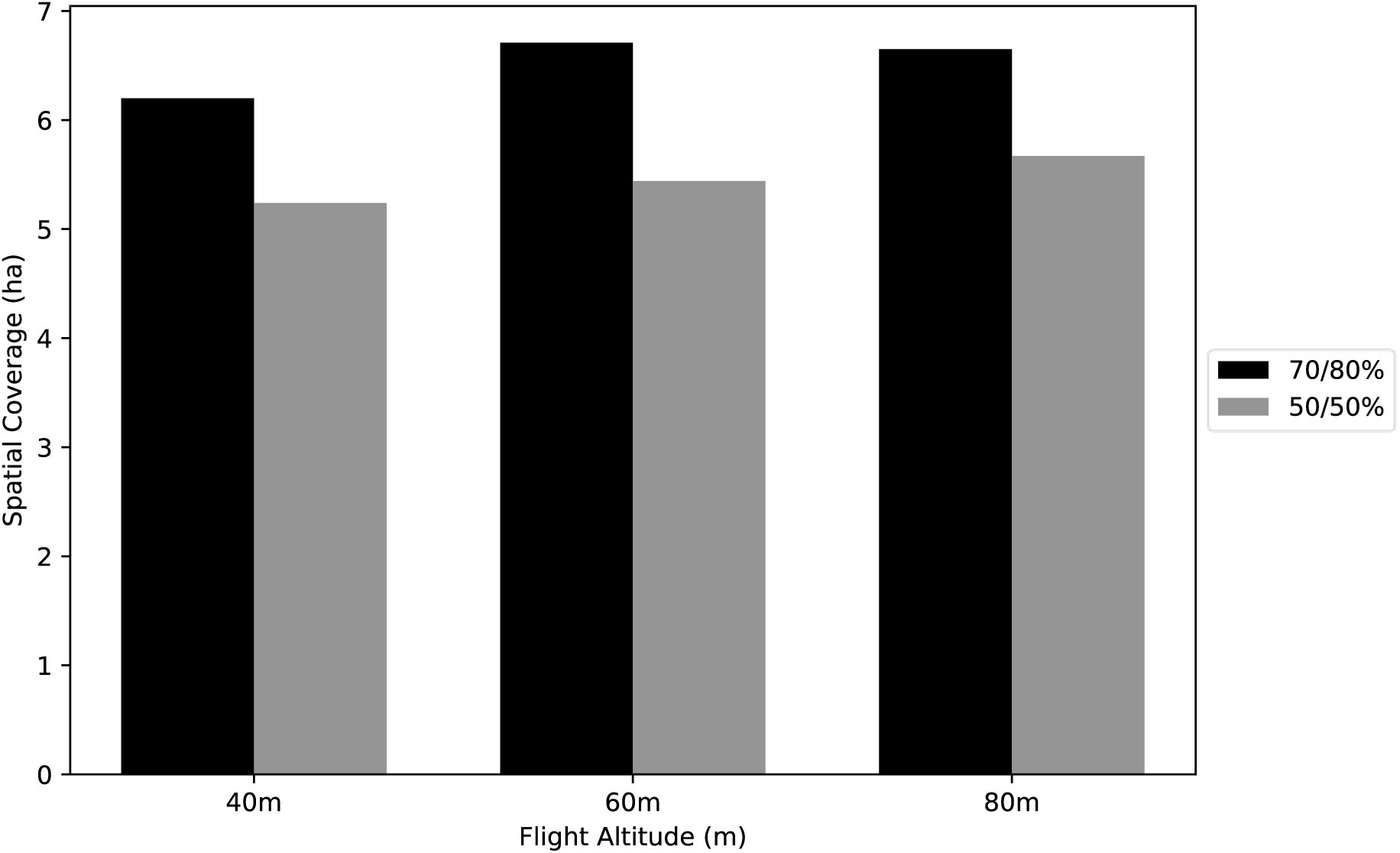
Spatial coverage of flights conducted at different altitudes, with overlap percentages indicated in the legend. Data for 20/20% overlap were excluded due to failure in orthomosaic generation resulting from insufficient image alignment. The chart presents data from Site A1 only.

+Table 5 shows that lower altitudes corresponded with smaller Ground Sampling Distances (GSD), indicating higher image resolution. The image resolution increases (GSD decreases) when the drone was operated at a lower altitude as shown in Figure 4.

**Figure 4.**
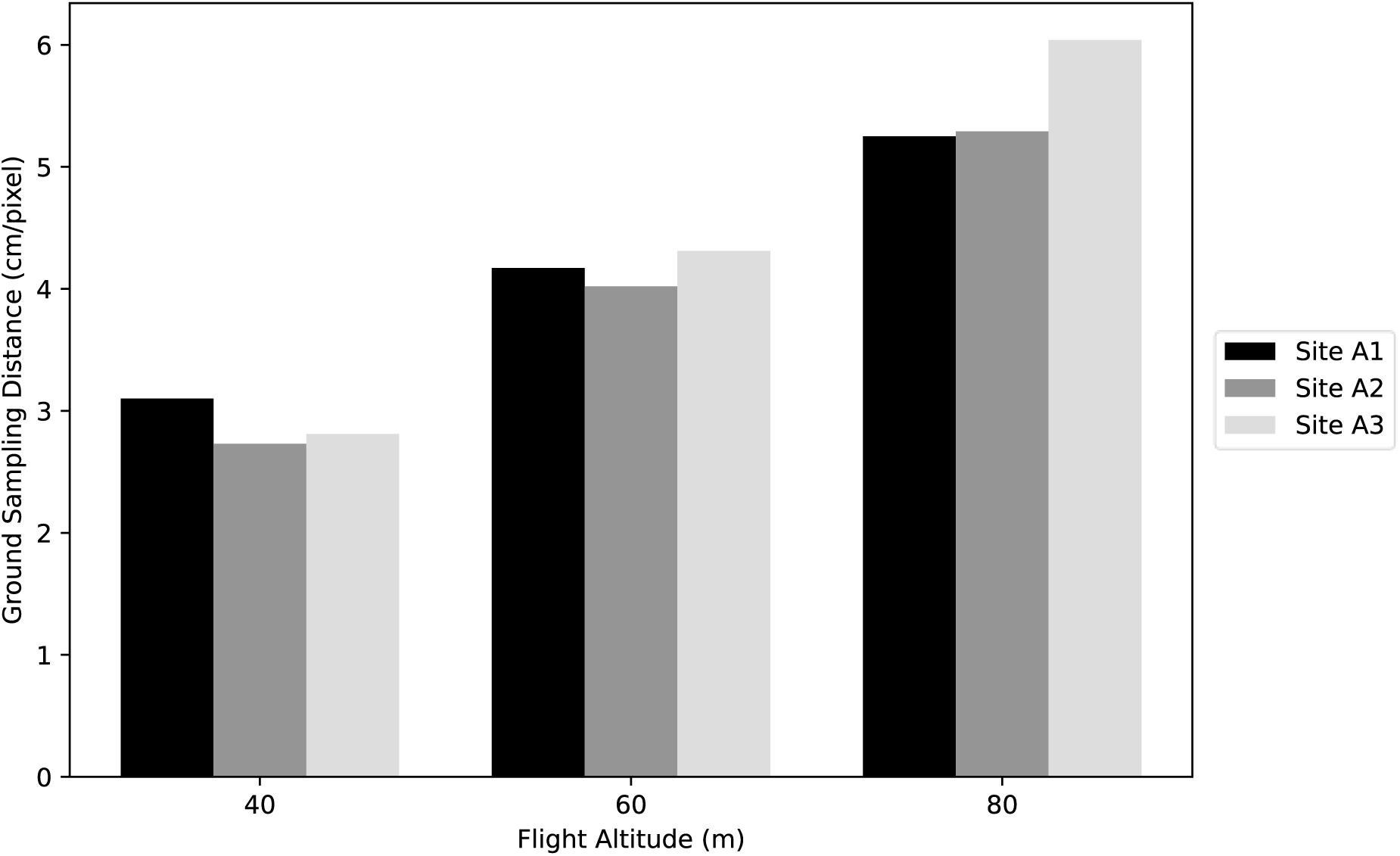
Ground sampling distance (GSD) in cm/pixel across three study sites (A1, A2, A3) at flight altitudes of 40 m, 60 m, and 80 m. All measurements were obtained from drone surveys with 70/80% image overlap.

**Table 5.** Ground sampling distance (cm/pixel) for the three study sites (A1, A2, A3) across different flight altitudes and image overlap percentages. Asterisks (*) indicate parameter combinations where orthomosaics could not be generated due to insufficient image alignment. Em dashes (—) denote incomplete data collection at Site A3.

| Site | Overlap % | Ground sampling distance (cm/pixel) |  |  |
| --- | --- | --- | --- | --- |
|  |  | 40m AGL | 60m AGL | 80m AGL |
| A1 | 70/80 | 3.1 | 4.17 | 5.25 |
|  | 50/50 | 2.73 | 4 | 5.43 |
|  | 20/20 | * | * | * |
| A2 | 70/80 | 2.73 | 4.02 | 5.29 |
|  | 50/50 | 2.8 | 3.89 | * |
|  | 20/20 | * | * | * |
| A3 | 70/80 | 2.81 | 4.31 | 6.04 |
|  | 50/50 | — | — | 5.34 |
|  | 20/20 | * | * | * |

##### Operational Efficiency

Operational efficiency was quantified by calculating area coverage (ha/min) for each viable parameter set (Table 6). This metric revealed that higher altitude flights with moderate overlap percentages achieved optimal efficiency. At Site A2, coverage efficiency increased nearly five-fold from 0.16 ha/min at 40 metres with 70/80% overlap to 0.80 ha/min at 80 metres. Similarly, Site A1 showed efficiency improvements from 0.21 ha/min to 0.95 ha/min across the same parameter range.

**Table 6.** Area coverage rates (ha/min) for each site across different flight altitudes and overlap percentages. Superscript plus signs (⁺) indicate the optimum flight parameter combination (60 m above ground level, 50/50% overlap) based on the balance between resolution and efficiency. Em dashes (—) indicate parameter combinations where orthomosaics could not be generated due to insufficient image alignment. Data for 20/20% overlap is excluded for the same reason.

| Site | Overlap % | Area coverage rate (ha/min) |  |  |
| --- | --- | --- | --- | --- |
|  |  | 40 m AGL | 60 m AGL | 80 m AGL |
| A1 | 70/80 | 0.21 | 0.48 | 0.95 |
|  | 50/50 | 0.29 | 0.68<sup>+</sup> | 1.42 |
| A2 | 70/80 | 0.16 | 0.41 | 0.8 |
|  | 50/50 | 0.25 | 0.69<sup>+</sup> | — |
| A3 | 70/80 | 0.15 | 0.48 | 0.99 |
|  | 50/50 | — | — | 1.31 |

However, these efficiency gains at 80 metres AGL must be weighed against two important constraints. First, successful orthomosaic generation at this altitude required higher overlap percentages (70/80%), as attempts with lower overlap often failed due to insufficient image alignment. Second, while flights operated at 80 metres AGL maximised area coverage efficiency, the resulting thermal resolution created challenges for macropod identification that will be discussed in Section 4.1.

### 3.2 Manual annotation

Manual annotation time varied with both image resolution and macropod density. At lower altitudes (40 metres AGL), higher resolution thermal images resulted in more detailed orthomosaics, allowing for clearer visualisation of individual macropods. However, this increased level of detail required more meticulous inspection during the annotation process, leading to longer annotation times per hectare (min/ha). For example, at Site A2 with 70/80% overlap, annotation took 1.72 min/ha at 40 metres AGL compared to just 1.01 min/ha at 80 metres AGL (Table 7).

**Table 7.** Time required for manual annotation (minutes per hectare) across different flight altitudes and image overlap percentages for each study site. Em dashes (—) indicate parameter combinations where orthomosaics were not generated due to insufficient image alignment.

| Site | Overlap % | Min/ha |  |  |
| --- | --- | --- | --- | --- |
|  |  | 40m AGL | 60 m AGL | 80 m AGL |
| A1 | 70/80 | 0.73 | 0.54 | 0.74 |
|  | 50/50 | 2.5 | 1.53 | 1.47 |
| A2 | 70/80 | 1.72 | 1.41 | 1.01 |
|  | 50/50 | 1.44 | 0.81 | — |
| A3 | 70/80 | 1.66 | 1.38 | 0.19 |
|  | 50/50 | — | — | 1.07 |

In contrast, flights at higher altitudes (80 metres AGL) generated coarser imagery but covered larger areas per flight. Although the reduced resolution made individual animals less distinct, the greater spatial coverage per orthomosaic generally resulted in lower annotation times per hectare. This is an inevitable trade-off between image clarity and annotation efficiency, where higher-resolution images offer greater confidence in identification but require more time to process, while lower-resolution images allow for faster annotation across broader areas, albeit with potential reductions in accuracy.

The rate of macropod detection per minute (macropods/min) varied with flight parameters and was influenced by factors such as macropod density and thermal contrast. Higher densities at Sites A2 and A3 often allowed for quicker identification due to frequent and prominent animal appearances in the imagery, particularly at 60 and 80 metres AGL where wider coverage facilitated efficient scanning. However, while detection rates were higher, the total time required for annotation was also longer for these denser image sets. In contrast, lower detection rates (e.g. at Site A1, 60 metres AGL, 70/80%, Table 8) reflected both fewer macropods and reduced thermal contrast, which made animals harder to identify. Annotation of the ‘warm’ orthomosaics was slower overall due to diminished thermal contrast, regardless of altitude or overlap.

**Table 8.** Detection rate of macropods (individuals per minute) during manual annotation of thermal orthomosaics, across varying altitudes and overlap percentages at three study sites. Em dashes (—) indicate parameter combinations where orthomosaics could not be generated due to insufficient image alignment.

| Site | Overlap % | Macropods/min |  |  |
| --- | --- | --- | --- | --- |
|  |  | 40m AGL | 60 m AGL | 80 m AGL |
| A1 | 70/80 | 19.91 | 8.77 | 15.42 |
|  | 50/50 | 15.6 | 23.01 | 21.88 |
| A2 | 70/80 | 16.61 | 31.99 | 32.82 |
|  | 50/50 | 25.12 | 53.05 | — |
| A3 | 70/80 | 17.08 | 22.96 | 16.89 |
|  | 50/50 | — | — | 23.08 |

### 3.3 Thresholding

Three orthomosaics, each captured at a different survey altitude with 70/80% overlap from the ‘cold’ dataset, are presented in this section. The ‘warm’ datasets were excluded from this analysis due to potential bias introduced by elevated residual ground heat, which compromised thermal contrast, as shown in Figure 5.

**Figure 5.**
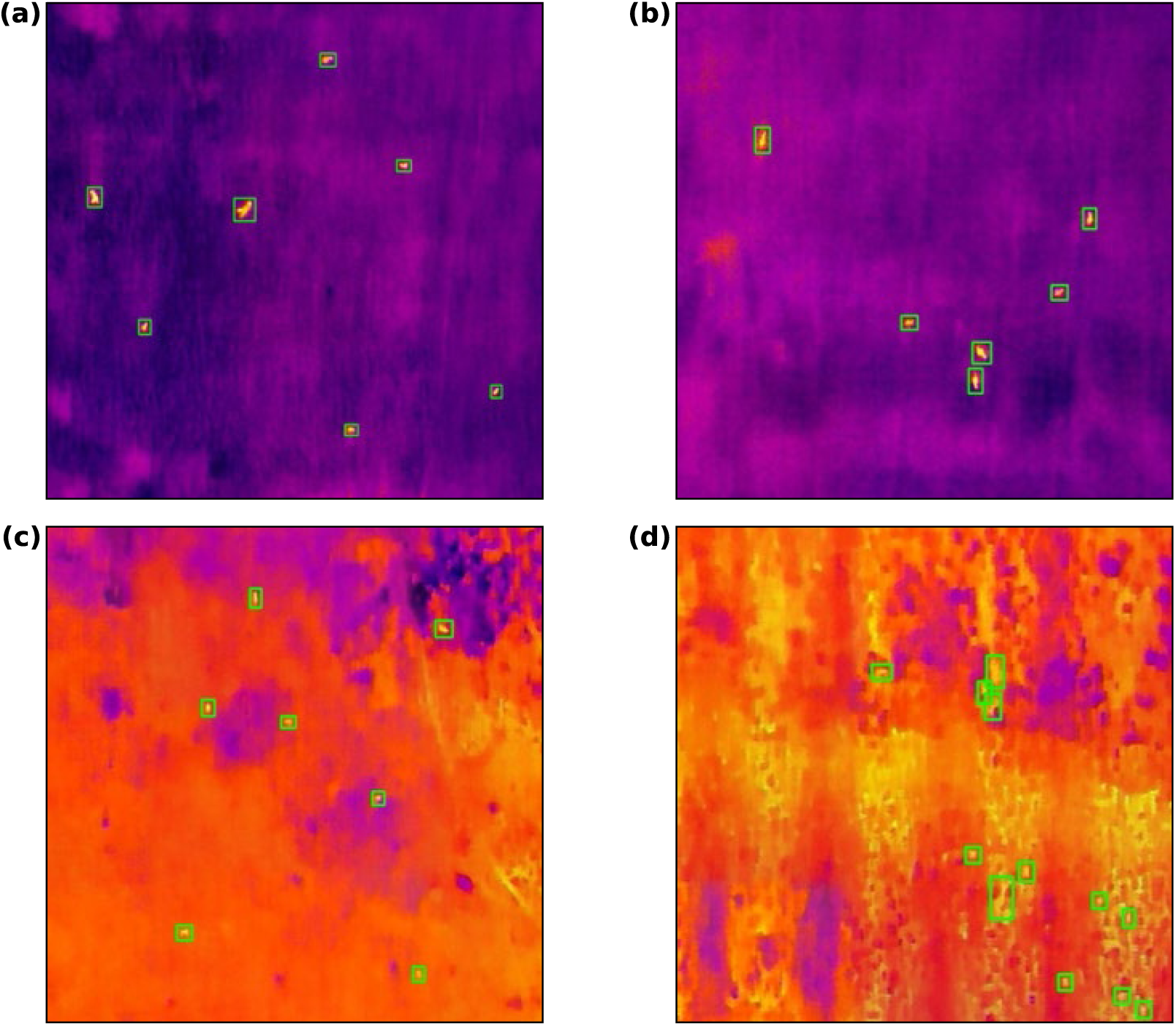
Examples of cropped sections of thermal orthomosaics from ‘cold’ sets (a, b) and ‘warm’ sets (c, d), with macropod detections shown as bounding boxes. Thresholding performs effectively when there is strong thermal contrast between the background and target species, as seen in (a) and (b). In (c), macropods remain detectable despite elevated background temperatures. In contrast, (d) demonstrates a detection failure case where similar thermal signatures between the animals and their surroundings result in false positives and inaccurate detection.

Manual annotation required more time than the automated thresholding approach (Table 9), but given the relatively small survey area, the workload remained manageable.

**Table 9.** Comparison of processing time (minutes) for manual macropod counts versus thresholding at three flight altitudes with 70/80% image overlap. Processing times are reported for the full orthomosaics. The 60 m above ground level (AGL) image set contained a higher number of macropods, resulting in longer processing times for both methods.

| Altitude (m) | Time taken (min) |  |
| --- | --- | --- |
|  | Thresholding | Manual |
| 40 | 0.5 | 4.52 |
| 60 | 7.78 | 30.23 |
| 80 | 0.1 | 3.02 |

The performance of the thresholding method was evaluated using the F1 score, calculated for each orthomosaic (Table 10). Full details on the performance metrics and the mathematical formulation of the F1 score are provided in Section 2.4.3.

**Table 10.** Comparison of macropod counts obtained via thresholding and manual annotation across different flight altitudes, along with the F1 score of the thresholding method. F1 scores indicate strong agreement between methods, particularly at 60 m and 80 m above ground level (AGL).

| Altitude (m) | Macropod count (thresholding) | Macropod count (manual) | F1 score (thresholding) |
| --- | --- | --- | --- |
| 40 | 91 | 89 | 0.91 |
| 60 | 943 | 921 | 0.96 |
| 80 | 47 | 51 | 0.96 |

## 4 Discussion

### 4.1 Flight parameters

In this study, optimal combinations of flight parameters for detecting macropods using thermal imagery under low-light conditions were identified as 60 metres AGL with 50/50% image overlap. Lower flight altitudes (e.g. 40 metres AGL) yielded higher-resolution imagery, which enabled clearer visualisation of macropod body shapes (i.e. outline of the macropod’s tail), helping to distinguish macropods from non-target species, such as brushtail possums (*Trichosurus vulpecula*), thereby supporting more confident detection of the target species. However, these gains in image quality came at the cost of reduced spatial coverage and longer flight times. For instance, 40 metres flights with 70/80% overlap took over two hours at the larger Site A2, necessitating multiple battery swaps and increasing logistical burdens.

In contrast, flights conducted at 80 metres AGL covered significantly larger areas in a shorter time. However, this operational efficiency was offset by a reduction in image resolution, which hindered the identification of individual animals (Figure 6), as reported in Brunton, Leon and Burnett (2020).. These findings align with previous studies showing that lower flight altitudes improve thermal resolution but reduce efficiency due to narrower fields of view and longer flight durations (Flores-De-Santiago *et al.* 2020; Rahman, Sitorus & Condro 2021; Pinel-Ramos *et al.* 2024). This trade-off was anticipated, given the known limitations of compact drone-mounted thermal sensors in balancing coverage and resolution. These outcomes underscore the need to align flight parameters with study goals, whether prioritising spatial coverage or reliable identification. In this study, despite the lower visual resolution at 80 metres AGL, F1 scores remained high across altitudes. This is because the thresholding algorithm used in this study relies on relative thermal intensity differences between macropods and the background rather than absolute image detail, such as the outline or shape of macropods. As noted in Section 3.3, only ‘cold’ orthomosaics, which provided high thermal contrast, were included in the analysis, ensuring reliable threshold-based detection. Therefore, although visual resolution decreased with altitude, as shown in Figure 6, F1 score continued to reflect high threshold-based detection performance (Table 10).

**Figure 6.**
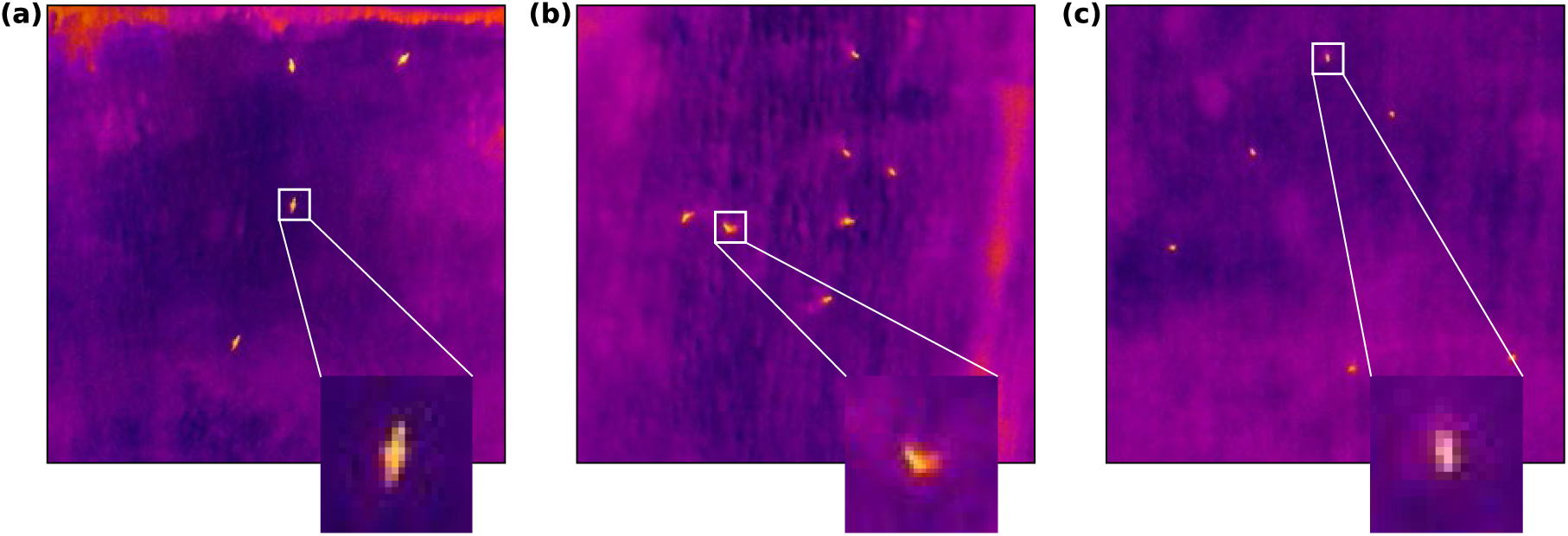
Examples of macropod detection at three flight altitudes: (a) 40 m above ground level (AGL), (b) 60 m AGL, and (c) 80 m AGL. Images demonstrate the decreasing visual resolution of macropods as altitude increases.

### 4.2 Habitat and environmental conditions

Although the study sites appear uniformly open, two environmental factors constrained thermal detection. First, micro-topographic features such as shallow undulations and remnant plough lines introduced partial occlusion that could physically obscure animals from the thermal sensor, particularly smaller individuals resting close to the ground. These features also create local variations in surface temperature, producing thermal “noise.” On warm nights, ridges and depressions in plough lines can appear brighter than the surrounding ground, either masking animal signatures or creating false positives in thermal imagery (Figure 7). Such small-scale variations illustrate that even minor terrain features can affect the detectability of animals. Terrain and vegetation characteristics have been shown to influence detection probability in thermal drone surveys (Blum *et al.* 2024), suggesting that similar features may affect thermal detection. Second, thermal background conditions influenced image clarity and contrast. Residual ground heat on warm nights narrowed the temperature gap between macropods and their surroundings, producing low-contrast images in which animal signatures merged with the background. Similar effects of elevated ground temperatures on reduced thermal detectability have been reported in other wildlife studies (Rahman *et al.* 2020; Lappin *et al.* 2024). Thus, openness alone did not ensure high detectability; prevailing thermal conditions proved equally critical. This underscores the need to factor both habitat and prevailing thermal background into nocturnal-flight planning.

**Figure 7.**
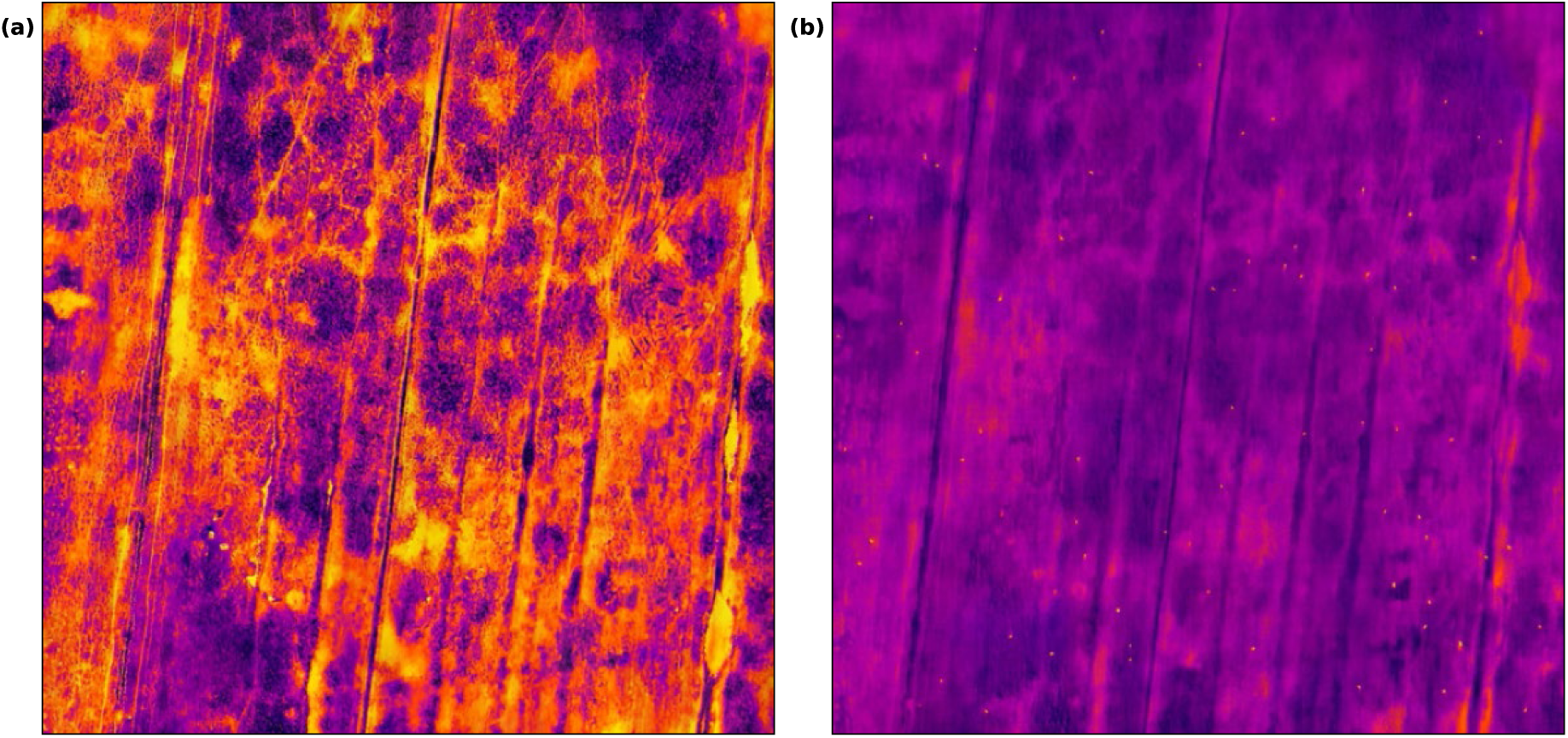
Cropped thermal orthomosaic showing micro-topographic features at the study sites. Panel (a) illustrates shallow plough lines and undulations, which create local variations in surface temperature and produce thermal “noise.” On warm nights, ridges and depressions in plough lines can appear brighter than the surrounding ground. Panel (b) shows the same location on a different night, highlighting how these features can reduce thermal contrast between animals and the surrounding ground, potentially masking animal signatures or creating false positives.

Weather, season, and survey timing also shape thermal image quality (Burke *et al.* 2019a). In forested habitats, morning flights often generate ‘thermal clutter’, as sun-lit tree trunks warm unevenly, whereas evening flights can yield clearer signatures once vegetation retains heat and the ground cools (Witczuk *et al.* 2018; Beaver *et al.* 2020). Conversely, early evening flights (1–4 hours after sunset) have been reported to produce unreliable thermal signatures, both in surveys of koalas (*Phascolarctos cinereus*) in open woodland and swamp forest habitats of New South Wales (Beranek *et al.* 2020) and primates in tropical forests of Panama (Kays *et al.* 2019). In this study, no clear within-night differences emerged; instead, detection varied mainly between survey days. All flights were flown between 0000 and 0700 h in early October—well before sunrise (∼0650 h)—so direct solar heating had little influence on surface temperatures. In addition, our open-grassland setting differs from the forested habitats examined in previous work, further emphasising that optimal survey windows are habitat-specific and must account for local thermal conditions.

Thermal contrast in this study was strongly influenced by temperature conditions leading up to and during the survey nights. Nights following hot, sunny days (e.g., 30^th^ September: solar exposure = 19.5 MJ m⁻²; minimum temperature = 11.8 °C) retained more ground heat, hence producing the lowest animal-background contrast recorded across all survey nights. By contrast, 1^st^ October—preceded by lower solar input and cooler air temperatures—yielded the clearest ‘cold’ orthomosaics. These results reinforce broader evidence that small temperature differentials, especially after warm days, degrade nocturnal thermal detectability (Rahman *et al.* 2020; Wijayanto, Condro & Rahman 2023; Lappin *et al.* 2024; Pinel-Ramos *et al.* 2024), suggesting that detectability is likely to vary both between nights and across seasons depending on recent weather conditions. Interestingly, the notably higher number of detections at 60 metres AGL compared to 40 metres and 80 metres AGL (Table 10) likely reflects differences in survey timing and spatial coverage. The 60 metres survey covered a larger area (21.51 ha versus 6.2 ha at 40 metres AGL and 15.78 ha at 80 metres AGL as shown in Table 4) and was conducted earlier in the night (0132 h), whereas the 40 metres and 80 metres surveys were conducted later (0545 h and 0620 h, respectively). These differences coincide with macropod activity patterns, as Bennett’s wallabies and Tasmanian pademelons are primarily nocturnal (Le Mar, McArthur & Statham 2003; Le Mar & McArthur 2005), while Forester kangaroos are both nocturnal and crepuscular (Lethbridge 2020). Because survey timing was not a primary focus of this study, the available data were insufficient to formally test this effect; however, future work could explicitly examine how night-time timing interacts with flight parameters and species activity to influence detection.

These temperature effects were mirrored in our image-processing results. Warm orthomosaics required lower threshold settings and still produced more false positives and misses, while ‘cold’ sets processed cleanly. Manual annotation faced similar challenges: reduced contrast caused macropods to blend into the background (Figure 8), obliging the annotator to rely on subtle cues and make more subjective judgements. Heat from the surrounding environment also obscured defining features such as body outline and movement, further complicating detection. Consequently, annotating warm datasets took longer and remained less reliable than annotating cooler, high-contrast imagery.

**Figure 8.**
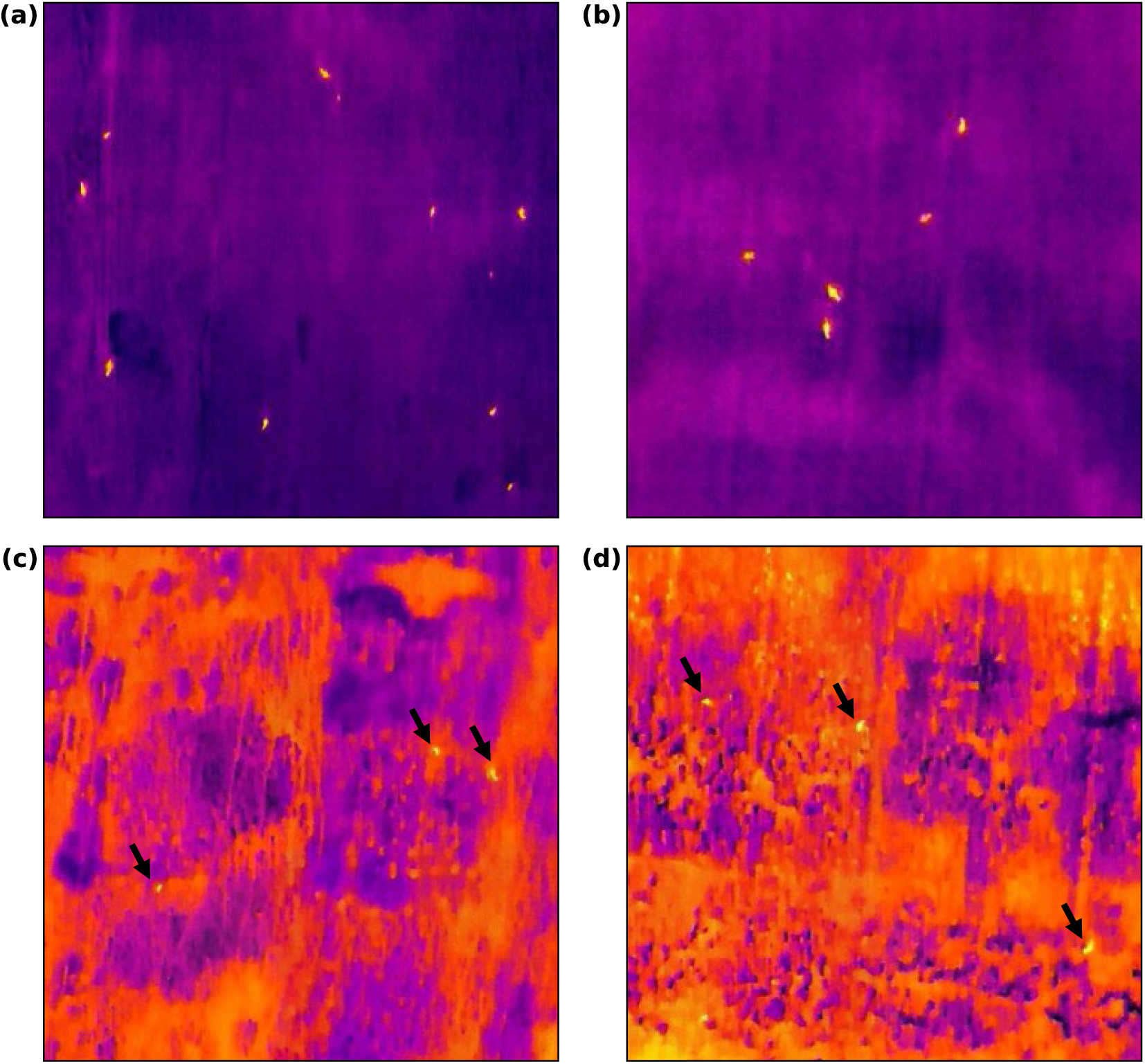
Examples of macropods captured on different survey dates and altitudes. Panels (a) and (b) show images taken on 1^st^ October at 60 m and 80 m above ground level (AGL), respectively, with high thermal contrast between the macropods and their surroundings. In contrast, panels (c) and (d), taken on 30^th^ September (at 80 m AGL) and 2^nd^ October (at 60 m AGL), respectively, display lower thermal contrast due to narrower temperature differentials. Macropods in panels (c) and (d) are indicated by black arrows.

### 4.3 Considerations in relation to ground-based methods

Nocturnal thermal drone surveys offer several advantages over traditional methods used for population monitoring, such as spotlighting, which remains common in Tasmania (NRET 2024). Spotlighting is constrained by a narrow field of view and observer variability. By contrast, thermal drones can deliver higher detection rates under suitable conditions. For example, Witt *et al.* (2020) reported that thermal drone surveys outperformed spotlighting in detecting koalas (*Phascolarctus cinereus*), likely due to complex vegetation structure and human observer error.

Spotlight surveys rely on a side or horizontal field of view, typically from vehicles or on foot (McGregor *et al.* 2021). This low-angle perspective means that any terrain undulations, vegetation, or other physical obstructions between the observer and the target animal can block visibility. Animals resting low to the ground or partially concealed by vegetation may be completely missed, particularly if they are not moving. In contrast, drones offer a top-down view that is much less affected by such obstructions; only dense overhead cover (e.g., tree canopies) will interfere with visibility. This aerial perspective reduces the risk of missing animals that are positioned behind shrubs or topographical features, which is an issue commonly encountered in side-profile spotlight surveys. In addition, drones can cover broader areas within a similar survey time, offering greater spatial efficiency compared to ground-based spotlighting (Witt *et al.* 2020).

Unlike spotlighting, which depends on eye-shine or movement for visual detection(Scott *et al.* 2005; Coulson, Snape & Cripps 2021; Hampshire & Miard 2024), thermal sensors passively capture heat signatures. This allows for detection of animals that are stationary, concealed, or facing away from the observer. Moreover, thermal imagery collected by drones can be archived, re-analysed, and independently validated, offering a level of repeatability and transparency that is not possible with observer-based ground counts.

### 4.4 Thermal drone imagery processing

Manual annotation remains the most accurate method for identifying animals in thermal imagery, particularly where detections are ambiguous or thermal contrast is low. However, manual review is time-intensive and becomes increasingly impractical as survey scales increase. In this study, preliminary results demonstrated that thresholding techniques can offer considerable time savings, less labour-intensive, and achieve reasonable performance (Table 10). Nonetheless, thresholding remains sensitive to background thermal variation as discussed in Section 4.2. Small fluctuations in environmental temperature or image noise can lead to false positives (e.g., hot rocks, patches of warm soil) or false negatives (e.g., animals with minimal temperature contrast).

Although manual annotation was optimal at this pilot scale, we suggest that thresholding could be integrated into a semi-automated workflow for larger surveys. For instance, thresholding could be used as a pre-screening tool in a semi-automated pipeline, particularly when paired with contextual filtering or integrated into machine learning approaches, such as CNNs. For small-scale or infrequent surveys, manual annotation remains feasible and preferred (Edney & Wood 2020). However, for broad-scale monitoring or high-frequency surveys, automation will become increasingly necessary (Beaver *et al.* 2020; Luz-Ricca *et al.* 2023).

Species-specific classification is challenging with low-resolution thermal imagery, particularly for small or partially occluded animals, which highlights the importance of selecting appropriate flight parameters. In this study, detection was performed at the macropod group level using thresholding, which was sufficient for the proof-of-concept objective of evaluating detection feasibility and workflow efficiency. Future research could explore more sophisticated machine learning approaches (e.g. CNNs), to enable species-level classification where image resolution and quality permit. Testing these frameworks across a broader range of environmental conditions and habitat types will help establish more robust and transferable workflows. Additionally, defining the threshold at which automation becomes necessary based on factors such as area, number of detections, or processing time, will be crucial for integrating these methods into long-term monitoring programs.

### 4.5 Limitations and challenges

A major limitation in the use of aerial thermal imagery is the difficulty in confidently identifying small or thermally indistinct animals. Large macropods such as Forester kangaroos can occasionally be identified from body size, outline and posture (Figure 9), but smaller macropods or non-target species (e.g., possums) often produce near-identical heat signatures—especially at higher altitudes. A juvenile wallaby and an adult pademelon, for instance, may be indistinguishable when no clear cues (limb movement, tail curvature) are visible (Figure 10). Similar identification constraints have been reported in several other studies (Goodenough *et al.* 2018; Burke *et al.* 2019b; Kays *et al.* 2019; Ivanova & Prosekov 2024). Some researchers have addressed this issue by using thermal video instead of still imagery (Goodenough *et al.* 2018; Kays *et al.* 2019), or by distinguishing species based on behavioural cues such as movement patterns, climbing behaviour, or group size (Burke *et al.* 2019b; Beranek *et al.* 2020; Ito, Fukue & Minami 2023). However, because large macropods in Tasmania often display similar behaviours and social structures, this approach is unlikely to enable species-level identification in this context. These challenges highlight the need for caution when interpreting thermal detections and suggest that incorporating additional data sources or conducting follow-up ground observations could improve accuracy for macropod detection.

**Figure 9.**
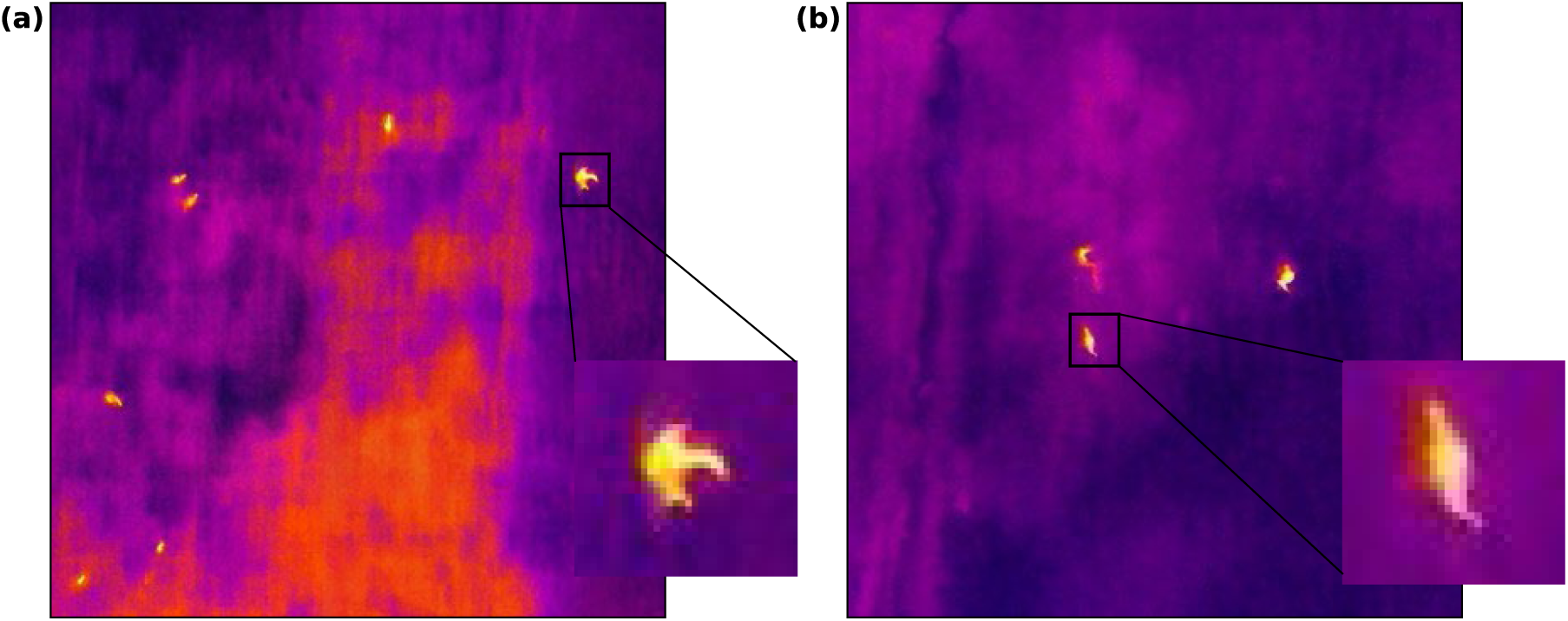
Examples of the distinct tail characteristics of a macropod observed in thermal imagery. Panel (a) shows the tail curvature captured at 60 m above ground level (AGL), while panel (b) illustrates the tail shape at 80 m AGL.

**Figure 10.**
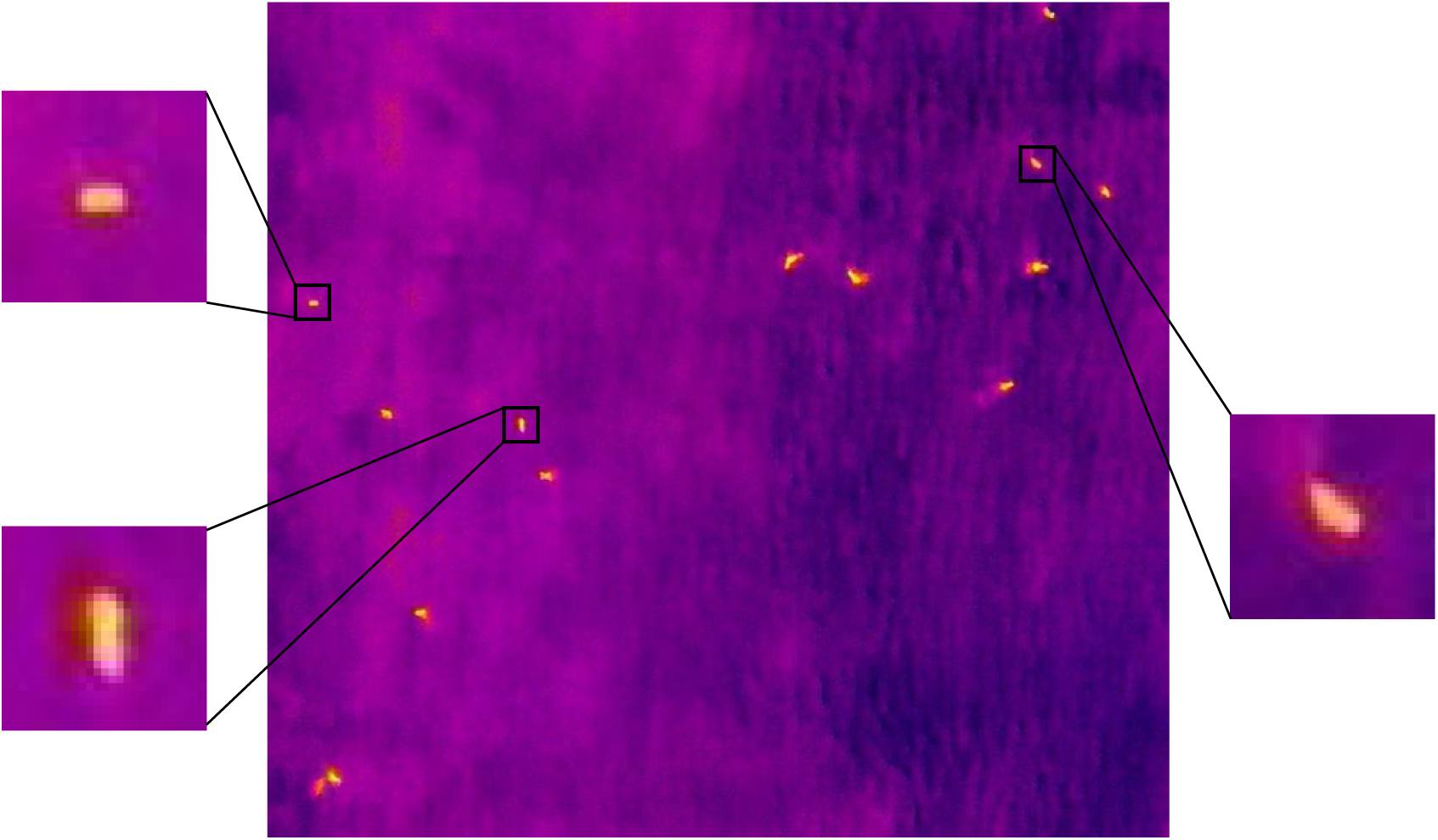
Example of macropods in thermal imagery where the lack of prominent identifying characteristics challenges detection and species-level identification. The image was captured at 60 m above ground level (AGL).

Environmental conditions also posed constraints on detection reliability. For instance, as highlighted in Section 4.2, low thermal contrast caused by residual ground heat during warm nights can result in thermal ‘blending’ between animals and their background. Survey timing is thus critical: cooler periods (e.g., early morning) may offer better thermal separation but must also align with the activity patterns of target species, which may be seasonally or behaviourally constrained.

Limited visibility at night increases the risk of collisions with obstacles such as power lines, trees, or buildings. The inability to maintain continuous visual line of sight (VLOS), as required under most drone regulations, further complicates navigation in low-light conditions (Linchant *et al.* 2015). This makes thorough pre-survey planning essential, including scouting the area during daylight hours, identifying potential hazards, and programming flight paths with appropriate safety buffers and altitude settings.

## 5 Conclusions

This study demonstrates that nocturnal drone surveys can detect macropods effectively in open Tasmanian landscapes. The findings suggest that flying at an altitude of 60 metres AGL with 50/50% image overlap offers an effective balance between spatial resolution and survey efficiency. This flight configuration produced high-resolution images suitable for macropod detection while ensuring sufficient overlap for orthomosaic reconstruction. In terms of image processing, manual annotation allowed for careful identification of macropods, although some detections required subjective judgement due to variable thermal contrast and potential confusion with background features. A simple thresholding approach offered considerable time savings and demonstrated a practical image analysis pipeline, but it is limited to detecting thermal intensity differences and does not provide species-level resolution. This workflow nonetheless provides a foundation for future studies to incorporate more sophisticated methods, such as CNNs, which could leverage high-resolution thermal imagery to enable species-level detection and identification. Such advancements would expand the applicability of thermal drone surveys for wildlife monitoring and could support species-specific management strategies where required. With continuing advances in thermal sensor technology, higher-resolution imagery may increasingly allow species-level detection even in challenging conditions.

Further research should investigate this approach in more complex habitats. Large-bodied macropods in Tasmania often occupy eucalypt forests and forest edges, where denser vegetation may reduce thermal visibility. In addition, seasonal variation should be explored to understand how changes in ambient temperature, vegetation cover, and animal behaviour affect thermal detectability throughout the year in Tasmania. Establishing reliable, low-disturbance methods for detecting nocturnal mammals has broader implications for wildlife management and conservation, particularly in improving the accuracy of population monitoring, informing management interventions, and supporting long-term ecological assessments in changing environments.

